# Gray Matter alterations in MS and CIS: a Coordinate based Meta-analysis and regression

**DOI:** 10.1101/2020.04.11.036954

**Authors:** Sonika Singh, Christopher Tench, Radu Tanasescu, Cris Constantinescu

**Affiliations:** Division of Clinical Neuroscience, University of Nottingham

**Keywords:** Coordinate based meta-analysis, voxel-based morphometry, grey matter atrophy, MS, CIS

## Abstract

The purpose of this coordinate based meta-analysis (CBMA) was to summarise the available evidence related to regional grey matter (GM) changes in patients with multiple sclerosis (MS) and clinically isolated syndrome (CIS). CBMA is a way to find the consistent results across multiple independent studies that are otherwise not easily comparable due to methodological differences. The coordinate based random effect size (CBRES) meta-analysis method utilizes the reported coordinates (foci of the clusters of GM loss) and Z score standardised by number of subjects, controlling type I error rate by false cluster discovery rate (FCDR). Thirty-four published articles reporting forty-five independent studies using voxel-based morphometry (VBM) for the assessment of GM atrophy between MS or CIS patients and healthy controls were identified from electronic databases. The primary meta-analysis identified clusters of spatially consistent cross-study reporting of GM atrophy; subgroup analyses and meta-regression were also performed. This meta-analysis demonstrates consistent areas of GM loss in MS or CIS, in the form of significant clusters. Some clusters also demonstrate correlation with disease duration.

## Introduction

Areas of inflammation, axonal loss, demyelination and gliosis, occurring throughout the brain and spinal cord, are the distinctive features of Multiple Sclerosis (MS). (Confavreux, Vukusic et al. 2000) Although MS has, in the past, been considered to be a condition affecting the white matter (WM) and the hyperintense lesions on T2 weighted images are the most important MRI characteristic for MS diagnostics, there is a limited association between lesion accrual and disability, reflecting pathological heterogeneity and consequent varied impact on clinical function and nerve conduction. Atrophy measures appear to be a more specific marker of MS pathology than lesion volumes(Miller, Barkhof et al. 2002), as demonstrated by the association of atrophy in the brain and spinal cord with increasing disability. (Losseff, Wang et al. 1996) In addition, progressive ventricular enlargement, another indicator of atrophy, has been shown to predate clinically definite MS in patients with clinically isolated syndrome (CIS) over a period of 1 year. (Dalton, Chard et al. 2004)

Atrophy of GM is already observed in the initial disease stages(Calabrese, Atzori et al. 2007) and an association has been observed with cognitive decline and physical disability(Messina and Patti 2014). The underlying mechanism for GM atrophy is still unknown. Several hypotheses have been postulated including primary GM damage involving neuronal loss, demyelination, reduced synapses, decreased oligodendrocytes as well as secondary damage involving axonal transection due to lesions. (Geurts and Barkhof 2008) An association has been demonstrated between GM loss and lesion load even in patients with short disease duration. (De Stefano, Matthews et al. 2003, Sailer, Fischl et al. 2003, Tedeschi, Lavorgna et al. 2005, Fisniku, Chard et al. 2008)

VBM can be used for the identification of localised regions of GM difference relative to a control group. Multiple VBM analyses of MS or CIS patients compared to healthy control groups have been published and significant changes interpreted as atrophy. When appropriately conducted, VBM has been shown to be robust against various processing steps with false positives randomly distributed about the brain. (Scarpazza, Tognin et al. 2015) However, studies often involve small sample sizes, and with lack of power comes increased chance that any observed effect is a false positive. (Button, Ioannidis et al. 2013) Furthermore, uncorrected p-values are commonly employed, inflating the false positive rates (Bennett, Wolford et al. 2009) and making the studies difficult to interpret.

A further complexity of VBM was highlighted by a study(Popescu, Schoonheim et al. 2016) comparing detectable GM changes by different software packages-FSL(Smith, Zhang et al. 2002), FreeSurfer(Dale, Fischl et al. 1999, Fischl, Sereno et al. 1999), SPM (Statistical Parametric Mapping Functional Imaging Laboratory, University College London, London, UK). The study examined agreement between these packages by using an MS cohort with a common disease type, a comparable disease duration, and controls that were matched. The study conducted by Popescu and colleagues(Popescu, Schoonheim et al. 2016) investigated the consistency in measurements of volume along with discrimination between patients and controls and, consistency of correlations with cognition. Consistency in volumes of GM regions was evaluated with Intraclass Correlation Coefficient (ICC) (consistency) for the lobar GM and deep GM (DGM) volume. The authors observed pronounced differences between the packages. The volume measurement consistency was substandard with a wide range of ICC values and, the lowest agreement structures included the amygdala, hippocampus, insula, and nucleus accumbens. Low ICC values signified potential reliability problems in relations with clinical variables as confirmed by inconstant results across methods. Nevertheless, good agreement was found between FreeSurfer and SPM in case of all cortical regions, with an ICC value of >0.7 for all, except left occipital lobe.

The deep GM (DGM) structure with high agreement in case of all methods was found to be the bilateral caudate nucleus. For bilateral thalamus, high agreement was observed between FSL and FreeSurfer. The DGM structures that include-bilateral thalamus, caudate nucleus, putamen, hippocampus and amygdala were found to have a significant difference between MS patients and controls in case of all methods.

Given the problems with single studies, there is potential for meta-analyses to reveal which of the observed effects are most likely to indicate common MS specific GM changes. In the absence of the original images, a coordinate based meta-analysis (CBMA) is possible using only the summary reports (coordinates and Z scores) tabulated in the large majority of VBM publications. The approach utilises the scientific necessity that any true and generalisable effect must be repeatably observable. The null hypothesis of CBMA is that the reported coordinates are randomly distributed, which reflects the spatial distribution of false positive observations in VBM. (Scarpazza, Tognin et al. 2015)

Coordinates that are concordant across multiple studies are an unlikely chance occurrence and so are critical of the null and declared significant.

The primary aim of this meta-analysis was to determine the locations of consistent GM changes in in MS and CIS patients by means of a coordinate based random effect size (CBRES) (Tench, Tanasescu et al. 2017) meta-analysis. This algorithm clusters the reported coordinates where there is spatial concordance, then performs conventional random effect meta-analysis, of the reported Z scores, in each cluster. The method has the advantage that censoring of effects due to the reporting of only summary results, rather than whole brain results, is dealt with statistically. Furthermore, CBRES offers principled control of the type 1 errors using the easily interpretable FCDR, which is based on the well-known FDR algorithm; FCDR is less conservative than the family-wise error rate (FWER) offered by other algorithms like activation likelihood estimation (ALE)(Laird, Fox et al. 2005) and multi-level kernel density analysis (MKDA)(Wager, Lindquist et al. 2009). Secondary analyses involving subgroup analysis and meta-regression are also performed using CBRES. In line with the suggestions of the preferred reporting items for systematic reviews and meta-analyses, the Royal Statistical Society, the coordinate data used in this analysis is provided as a supplement for validation purposes.

## Methods

### Search strategies

A literature search was conducted using PubMed - with the following search term combinations- (“multiple sclerosis”[All Fields] OR “ms”[All Fields] OR CIS[All Fields] OR “clinically isolated syndrome”[All Fields]) AND (“voxel based morphometry”[All Fields] OR VBM[All Fields]) AND (“atrophy”[MeSH Terms] OR “atrophy”[All Fields]) AND (“grey matter”[All Fields] OR “gray matter”[All Fields] OR GM[All Fields]), Web of science, using the following search terms-TS=(“multiple sclerosis” OR MS OR CIS OR “clinically isolated syndrome”) AND TS=(“voxel based morphometry” OR VBM) AND TS=(atrophy) AND TS=(“grey matter” OR “gray matter” OR GM) and Science direct, using the following search terms-TITLE-ABSTR-KEY(“multiple sclerosis” OR MS OR “clinically isolated syndrome” OR CIS) and TITLE-ABSTR-KEY(“voxel based morphometry” OR VBM) and TITLE-ABSTR-KEY(“grey matter” OR “gray matter” OR GM) and TITLE-ABSTR-KEY(atrophy). The studies included in our meta-analysis were conducted between 2006 and 2017.

### Study selection

The articles have been reviewed on the basis of stringent inclusion and exclusion criteria. A study was considered for inclusion if it (a) involved participants with MS or CIS (Table 1) (b) compared patients to healthy controls (c) performed whole brain VBM for assessing GM atrophy (d) reported coordinates for GM volume changes in either Talairach (Talairach 1988) or Montreal Neurological Institute (MNI) reference space.

**Table 1:**
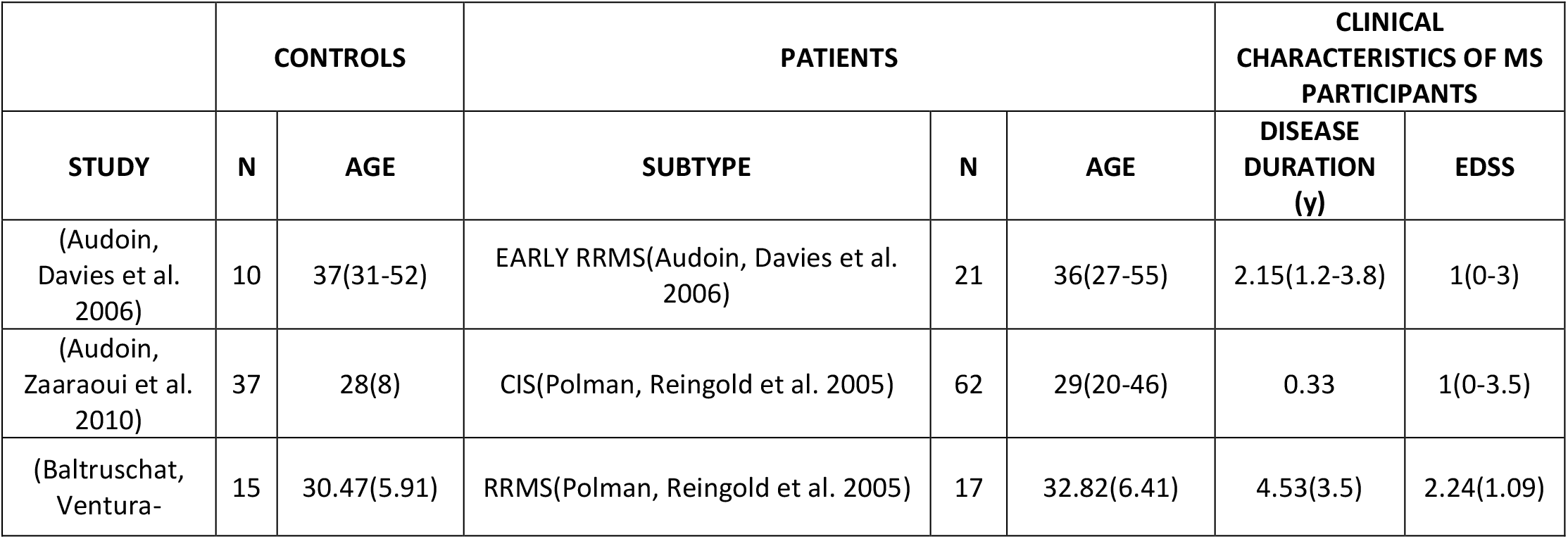
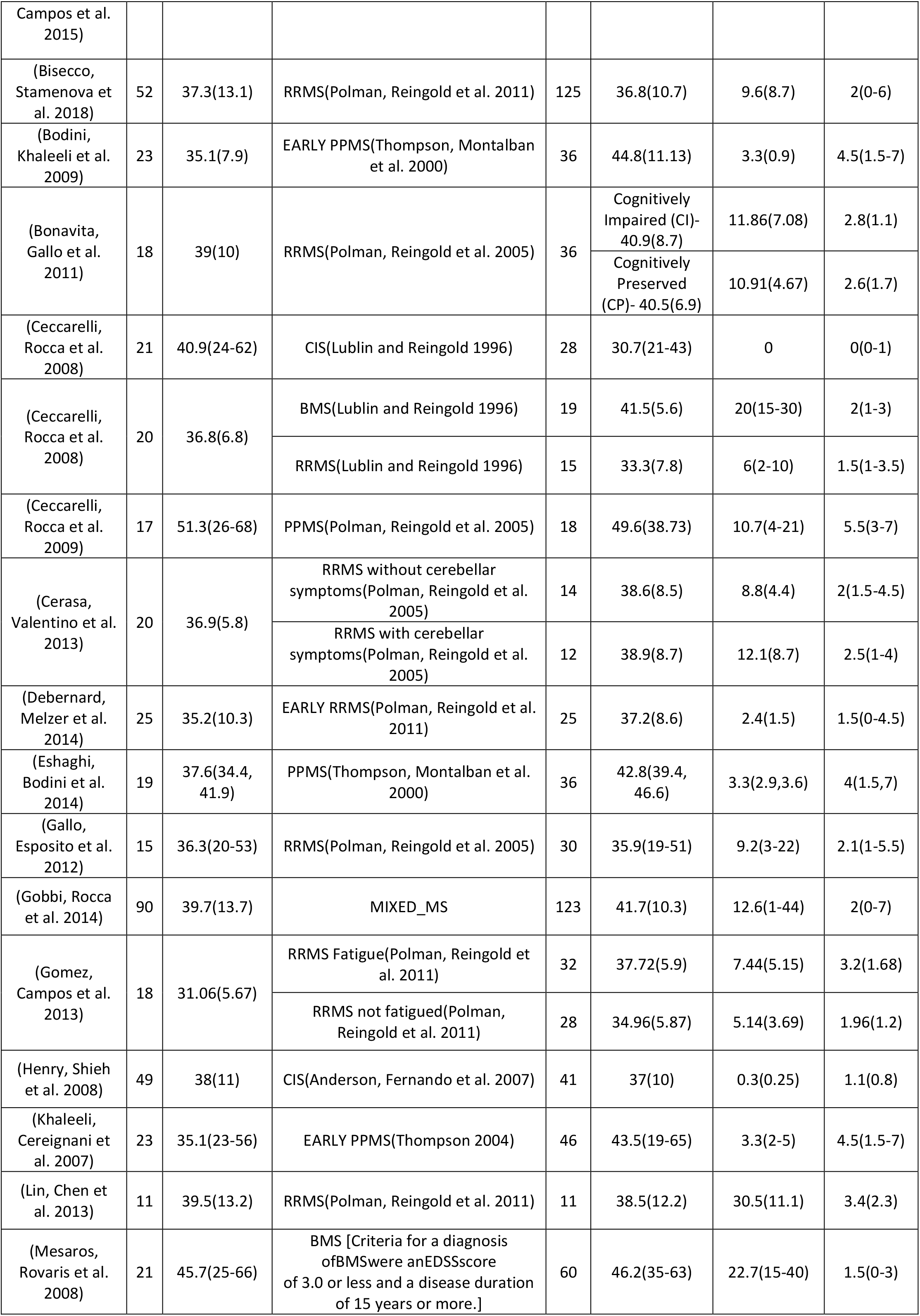
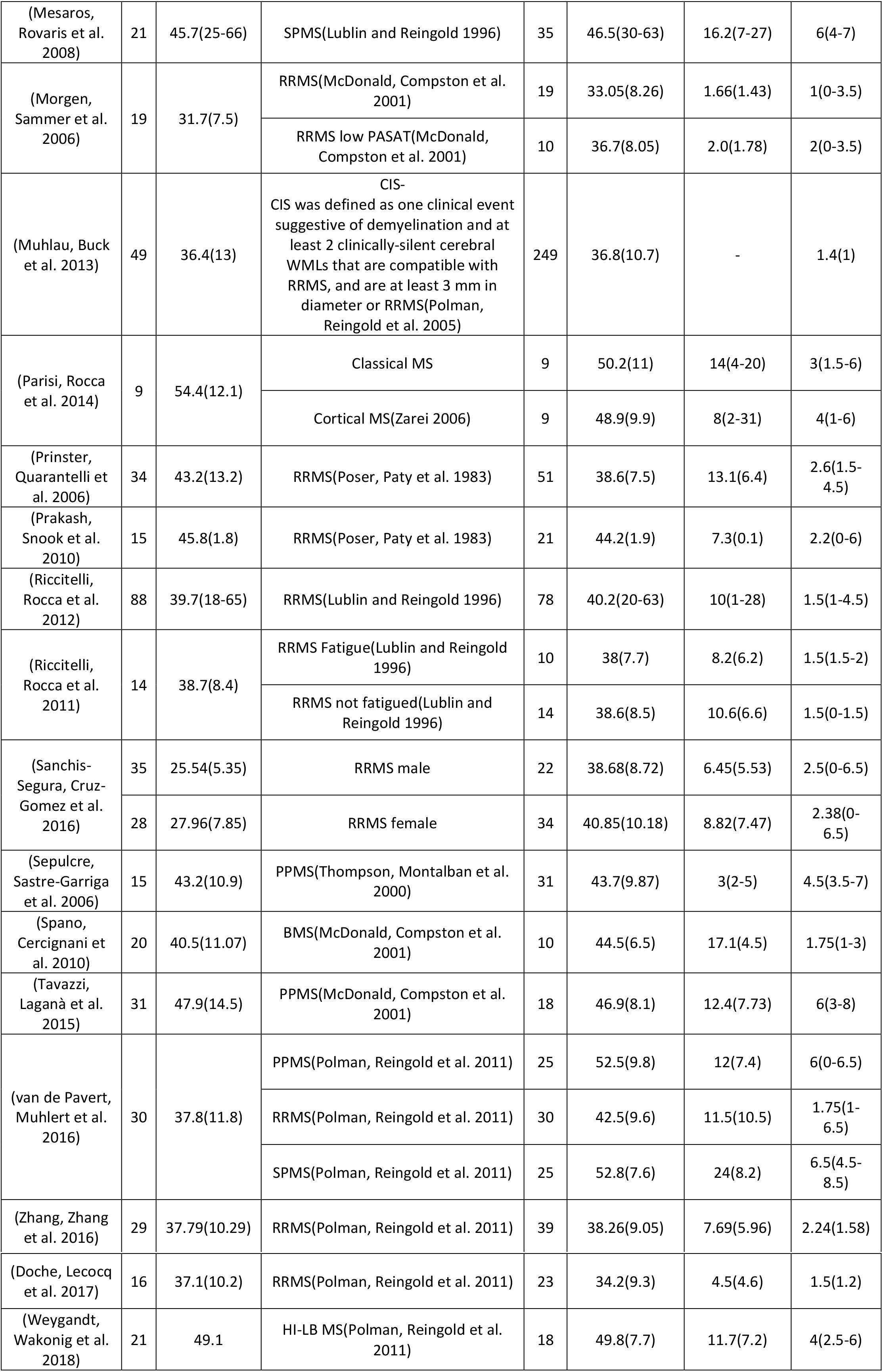
Demographics and clinical characteristics of datasets for studies included in the meta-analysis.

Exclusions were made due to unreported coordinates or unavailable full text. Two independent researchers assessed these criteria of the individual studies and the MNI or Talairach coordinates. Any disagreements between the researchers were agreed on by consulting and discussing the respective original article.

### Study properties

For each included study the following information is required for CBRES analysis: the censoring threshold i.e. the smallest Z value the study considered as significant, the reported coordinates, and either the Z score, estimated degrees of freedom and t statistic, or uncorrected p-value; t-statistics and uncorrected p-values are converted automatically to Z scores. Where no censoring threshold is reported, CBRES takes the lowest reported Z score as the threshold.

### Coordinate based random effect size Meta-analysis

All CBRES analysis is performed using ClusterZ, a free to use application that is part of NeuRoi (https://www.nottingham.ac.uk/research/groups/clinicalneurology/neuroi.aspx).

Details about the algorithms incorporated into ClusterZ are presented in (Tench, Tanasescu et al. 2017), but a brief overview is provided here. In CBRES a clustering algorithm (DBSCAN)(Sander, Ester et al. 1998)- (density-based spatial clustering of applications with noise) is used to determine where the coordinates reported by multiple independent studies are spatially concordant (clustered). To achieve this a clustering distance (CD) is needed; coordinates must fall within this distance of each other to form clusters. The CD is analogous to the full width half max (FWHM) in kernel based CBMA methods such as ALE and MKDA, but importantly it automatically adapts to the data to prevent increasing false positives as the number of studies increases. (Tench, Tanasescu et al. 2014) Once clusters are formed the reported Z scores are converted to standardised effect sizes by dividing by the square root of the number of subjects. Finally, a conventional random effect meta-analysis of these effect sizes is performed in each cluster. Where a study does not report a coordinate within a cluster, or where no effect sizes are reported by a study, the contribution of the study to the cluster is estimated using the censoring threshold. (Tench, Tanasescu et al. 2017)

The significant results of the CBRES meta-analysis are clusters of reported coordinates where the estimated effect size is statistically different to zero after controlling the false cluster discovery rate. This method of type 1 error control is based on FDR(Benjamini and Hochberg 1995), and works at the cluster level. The interpretation is that the expected proportion of clusters incorrectly declared significant (type 1 error) is controlled at a user specified level (usually 5%). These clusters indicate both spatial and effect size concordance across the multiple studies. Such concordance is an unlikely chance event, suggesting that atrophy at the location of the clusters is a consistent feature of MS. A feature of CBRES is that the clusters declared significant can be found as a function of FCDR. This means that the next most significant clusters are flagged, and any that just miss the threshold for significance can be explored.

ClusterZ also performs other types of analyses. A sub analysis can be performed on subgroups of studies. This estimates a subgroup specific effect size in each of the clusters found significant during the full analysis (using all studies); this is useful since clusters may not be significant if the subgroup is small, yet the effect size might be of interest.

Furthermore, the use of standardised effect sizes makes meta-regression possible. This analysis looks for significant correlation between a specified covariate and the standardised effect size in each cluster. Covariates can be continuous, such as age, or dichotomous to allow comparison of two independent groups. Meta-regression can be performed independently of the meta-analysis, or post-hoc only within clusters found to be significant during the meta-analysis.

### Experimental Procedure

Multiple experiments reported on the same subjects were pooled into single independent experiments. This prevents correlated results generated by the same subjects inducing apparent concordance that is not due to a generalizable MS process. (Turkeltaub, Eickhoff et al. 2012)

All planned analyses were performed controlling the FCDR at 0.05. For each the next most significant clusters were explored, and reported, to make sure that none had just been missed at this threshold.

#### Main analysis

The main meta-analysis was performed using all studies meeting the inclusion criteria.

#### Subanalyses

Subanalyses for CIS, benign MS (BMS) {30}, Relapsing Remitting MS (RRMS), Primary Progressive MS (PPMS) and Secondary Progressive MS (SPMS) studies were. This analysis estimates effects of just the respective subgroup within significant clusters discovered using all studies.

#### Meta-regression

Post hoc regression analyses were performed for covariates that might influence the grey matter volume: mean age (years), MS disease duration (years; excluding CIS studies with no MS disease duration), MSFC, and EDSS (all studies and including RRMS studies only).

## Results

### Included studies and sample characteristics

The literature search yielded 237 potential studies from which 34 met the inclusion criteria (Figure 1). The 34 included research papers reported 44 whole brain VBM experiments comparing MS subtypes and controls. (Table 1)

**Figure 1:**
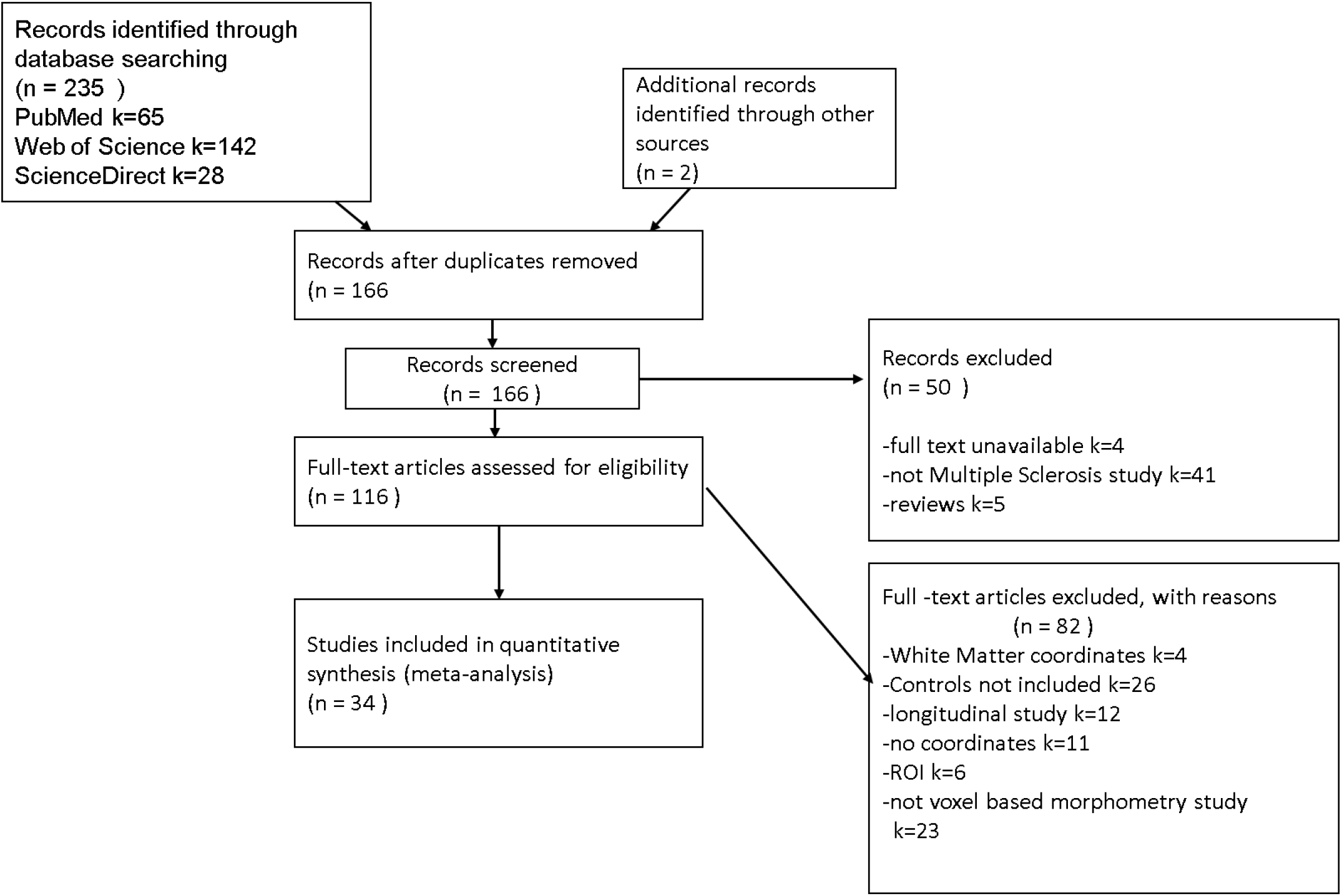
PRISMA flowchart(Liberati, Altman et al. 2009) showing the inclusion of studies in the meta-analysis and reasons for exclusion of studies.

The studies included in our meta-analysis were conducted between 2006 and 2017 and involved a total of 1561 patients and 1182 controls. The mean age of the patients was 40.49 years (SD = 5.64). The patients and controls were matched for age and gender in 21 out of 44 included studies. They were also matched for age only, in 3 studies, and males and females have been matched for age in 2 studies. The number of controls varied in size from 9 to 90 and patients varied from 9 to 249. All included studies reported the mean or median age, EDSS and disease duration. The mean (standard deviation (SD)) EDSS was calculated to be 2.71 (1.54). The mean disease duration was 9.26 years (6.51). One study including CIS or RRMS participants by Muhlau(Muhlau, Buck et al. 2013) did not report the disease duration.

Across the studies, the minimum duration of disease, excluding the CIS studies, was 1.66 years and maximum was 30.50 years.

### Primary meta-analysis

Analysis included 44 experiments and found 8 significant clusters. The location of significant clusters involved basal ganglia and cortical regions. The effect sizes of the clusters found using all studies are given in Table 3. The significant clusters and forest plots are shown in Figure 2 and Figure 3. The first non-significant cluster was at a FCDR of 0.12.

**Table 3:**
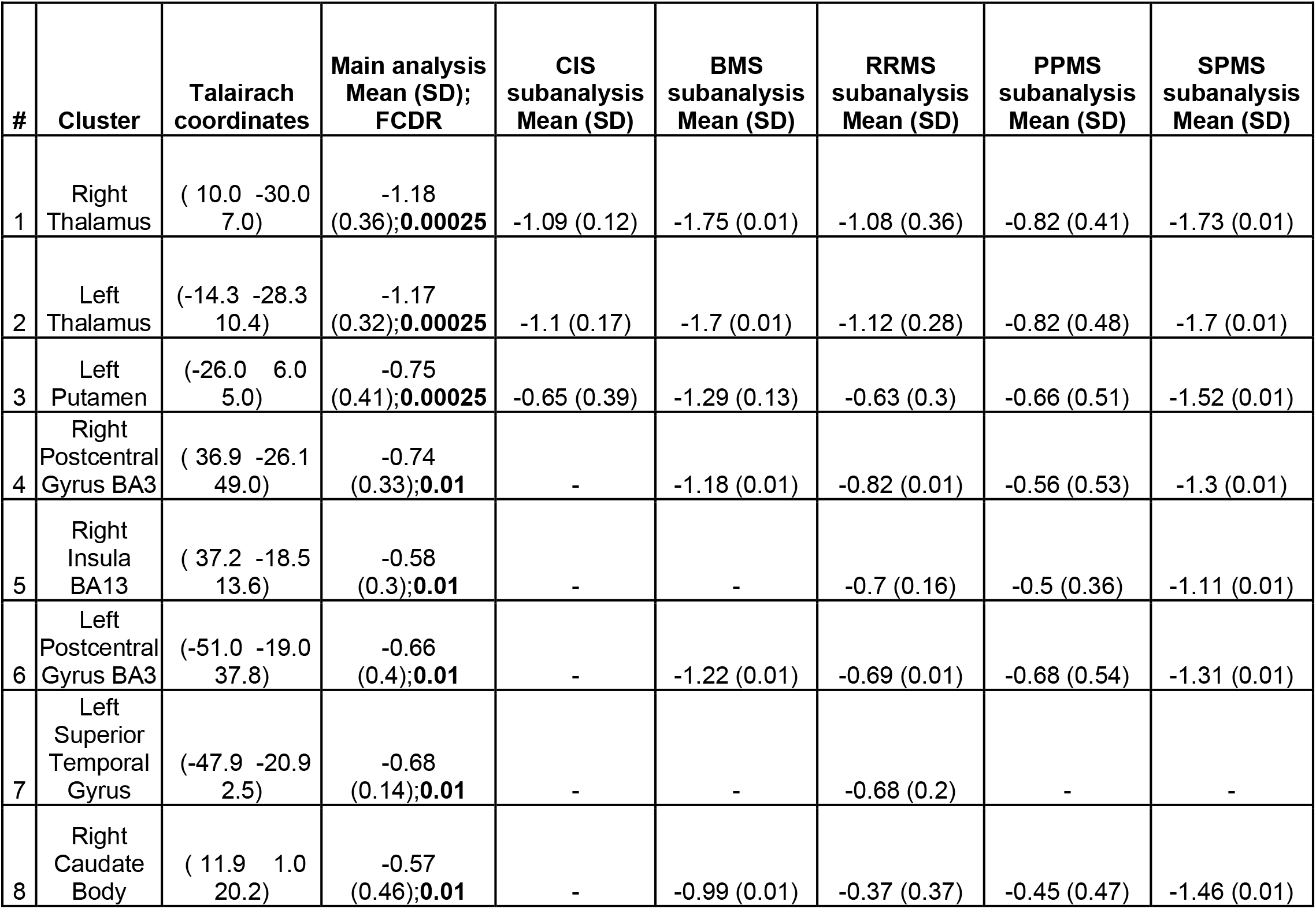
Table shows significant clusters for main analysis and estimated effects from the subanalyses. The column ‘main analysis’ shows effect size, standard deviation and p_value for each significant cluster. The following columns consist of effect size ± standard deviation for the subanalyses; - signifies no contribution of the subgroup to the cluster. (BA = Brodmann Area)

| # | Cluster | Talairach coordinates | Main analysis Mean (SD); FCDR | CIS subanalysis Mean (SD) | BMS subanalysis Mean (SD) | RRMS subanalysis Mean (SD) | PPMS subanalysis Mean (SD) | SPMS subanalysis Mean (SD) |
| --- | --- | --- | --- | --- | --- | --- | --- | --- |
| 1 | Right Thalamus | ( 10.0 -30.0 7.0) | -1.18 (0.36);**0.00025** | -1.09 (0.12) | -1.75 (0.01) | -1.08 (0.36) | -0.82 (0.41) | -1.73 (0.01) |
| 2 | Left Thalamus | (-14.3 -28.3 10.4) | -1.17 (0.32);**0.00025** | -1.1 (0.17) | -1.7 (0.01) | -1.12 (0.28) | -0.82 (0.48) | -1.7 (0.01) |
| 3 | Left Putamen | (-26.0 6.0 5.0) | -0.75 (0.41);**0.00025** | -0.65 (0.39) | -1.29 (0.13) | -0.63 (0.3) | -0.66 (0.51) | -1.52 (0.01) |
| 4 | Right Postcentral Gyrus BA3 | ( 36.9 -26.1 49.0) | -0.74 (0.33);**0.01** | - | -1.18 (0.01) | -0.82 (0.01) | -0.56 (0.53) | -1.3 (0.01) |
| 5 | Right Insula BA13 | ( 37.2 -18.5 13.6) | -0.58 (0.3);**0.01** | - | - | -0.7 (0.16) | -0.5 (0.36) | -1.11 (0.01) |
| 6 | Left Postcentral Gyrus BA3 | (-51.0 -19.0 37.8) | -0.66 (0.4);**0.01** | - | -1.22 (0.01) | -0.69 (0.01) | -0.68 (0.54) | -1.31 (0.01) |
| 7 | Left Superior Temporal Gyrus | (-47.9 -20.9 2.5) | -0.68 (0.14);**0.01** | - | - | -0.68 (0.2) | - | - |
| 8 | Right Caudate Body | ( 11.9 1.0 20.2) | -0.57 (0.46);**0.01** | - | -0.99 (0.01) | -0.37 (0.37) | -0.45 (0.47) | -1.46 (0.01) |

**Figure 2:**
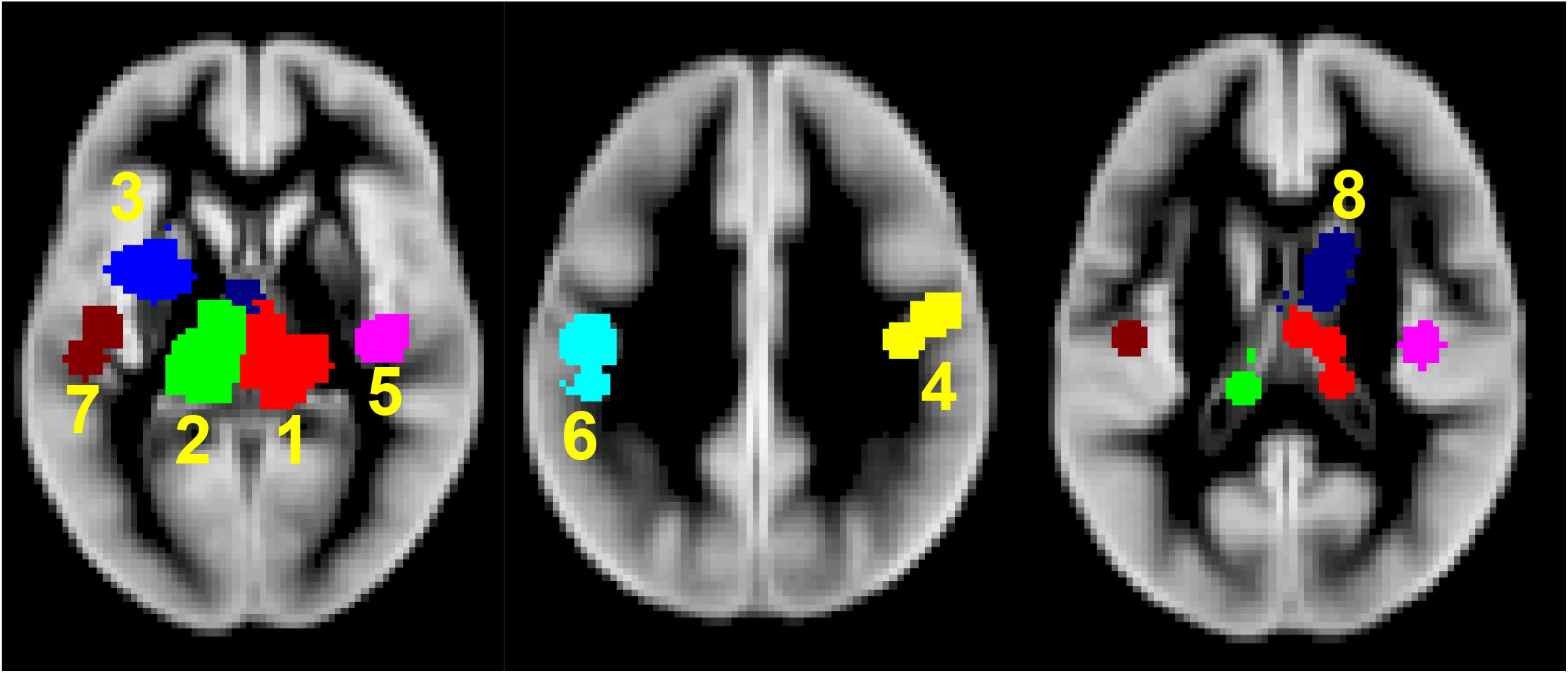
Significant clusters of GM atrophy reported by the 44 VBM experiments.1-Right Thalamus 2-Left Thalamus 3-Left Putamen 4-Right BA3 5-Right BA13 6-Left BA3 7-Left Superior Temporal Gyrus 8-Right Caudate Body (BA = Brodmann Area)

**Figure 3:**
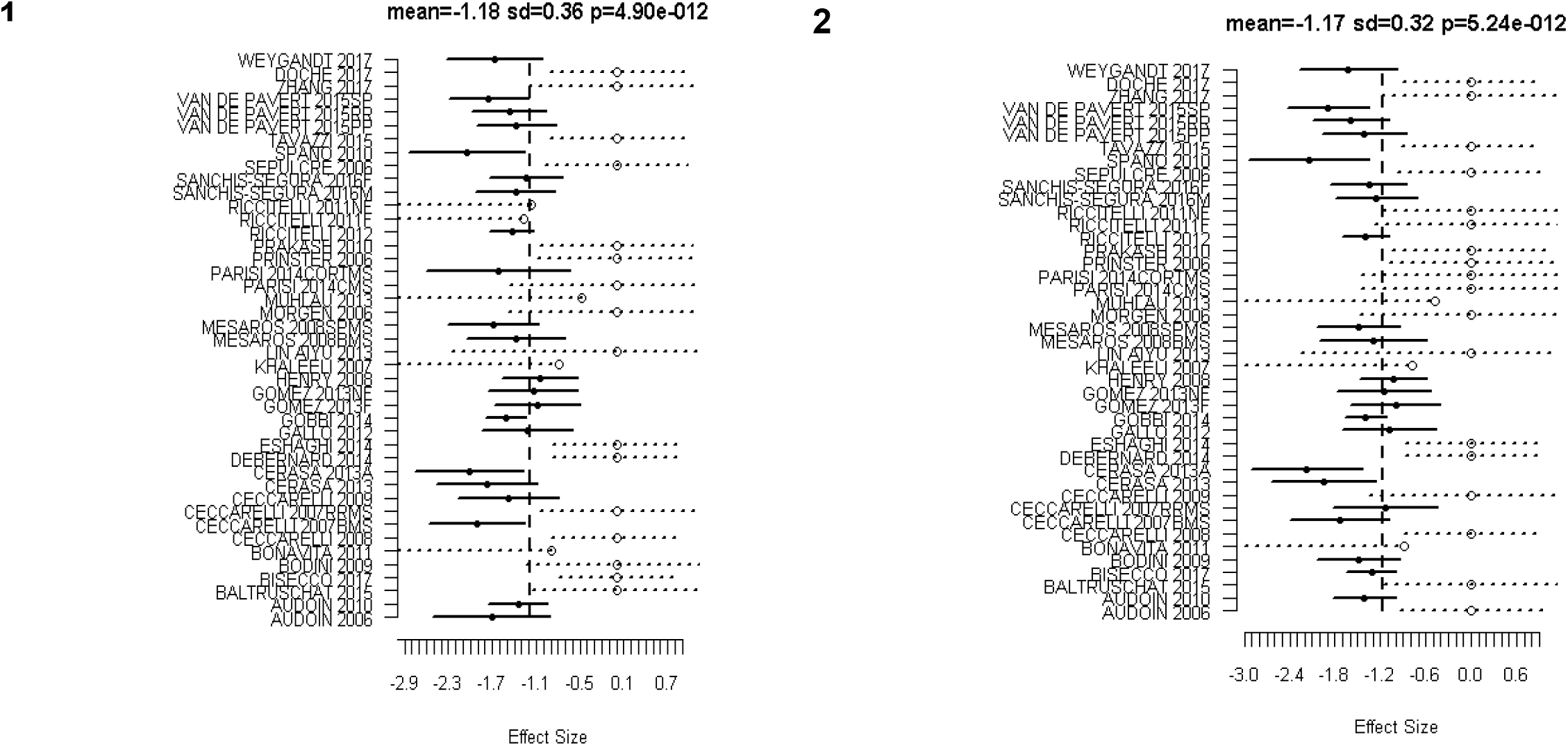
Forest Plots for the significant clusters (1 and 2) of GM atrophy reported by the 44 VBM experiments. Markers with solid circle indicate the effect size reported by the study in the respective cluster. The confidence intervals are depicted as solid horizontal lines spanning ± 1.96 times the within study standard deviation of the effect size. Censored values are depicted by open circle markers and the intervals by dashed lines (⋯o⋯).

#### Subanalyses

Results from the subanalysis are shown in table 3. It shows estimated effects sizes considering only the respective subgroup within each of the clusters from the primary meta-analysis. It is apparent that the estimated effect has the largest magnitude in the SPMS group.

### Metaregression

#### Age

ClusterZ found no clusters with age as a covariate. The first non-significant cluster was at FCDR 0.2.

#### MS Disease Duration

Two clusters with significant correlation between MS disease duration were found and reported; table 3 (figure 2, Cluster 1 and Cluster 2). The first non-significant cluster was found to be at FCDR 0.07 that is, just beyond the threshold of 0.05 for significance.

**Table 3:** Significant clusters for MS Disease Duration Meta Regression in all studies (excluding 4 CIS studies). Cov = Regression coefficient; Cov refers to change in effect size for every 1 year of disease duration.

| Cluster number | Talairach coordinates | p-value | Cov | Percentage of mean effect per year |
| --- | --- | --- | --- | --- |
| 1 (Right Thalamus) | ( 10.0 -30.0 7.0) | 0.01 | -0.06 | 5 |
| 2 (Left Thalamus) | (-14.3 -28.3 10.4) | 0.01 | -0.05 | 4 |

#### MSFC

Regression analysis could not be performed because only 9 out of 44 studies have given MSFC for MS patients.

#### EDSS

ClusterZ found no significant clusters. The first non-significant cluster was found to be at FCDR 0.30.

#### EDSS (RRMS only)

ClusterZ found no significant clusters. The first non-significant cluster was found to be at FCDR 0.25.

### Subgroup analysis

Planned comparison of effect sizes between patient subgroups was limited due to small numbers of studies in each group. In particular, there were only two SPMS studies, so no comparisons were performed with this subgroup. For completeness, however, no differences were found when comparing RRMS and PPMS (first cluster was found to be at FCDR 0.5), RRMS and BMS (first cluster was found to be at FCDR 1), CIS and RRMS (first cluster was found to be at FCDR 1), PPMS and BMS (first cluster was found to be at FCDR 0.5).

## Discussion

A commonly held view of MS, previously, involved the picture of a multifocal and multi-phasic immune-mediated WM inflammatory demyelinating disorder, the suppression of which is the basis of progress in DMTs to date. Nevertheless, it is quite clear as of now that GM is extensively involved as well. The literature makes it clear that GM pathology exists in early RRMS increasing with time. Brain atrophy in MS, as monitored during life by MRI, possibly reflects neuro-axonal loss (although other factors that can affect brain tissue volumes should be borne in mind, especially when assessing short-term changes).

### GM atrophy in MS: a pathological stance

GM tissue damage is an important pathological process in MS that underlies neurological disability. (Fisher, Lee et al. 2008) The loss of volume is the result of many dynamic processes, with a balance between destructive and reparative mechanisms with interaction among neurons, oligodendrocytes, axons, microglia, astrocytes, inflammatory cells, endothelial cells and water distribution. (Koskimäki, Bernard et al. 2018)

Previous literature shows differences in GM atrophy between MS subtypes involving a selective myriad of brain regions along with an increased extent of atrophy in common regions, such as the thalamus, in the progressive phase of the disease. (Gilmore, Donaldson et al. 2009) The reason of GM pathogenesis is not quite clear, however many suggestions regarding relevant pathological processes have been put forward, comprising meningeal inflammation and the secretion of myelinotoxic factors, retrograde myelino-axonal degradation, virtual hypoxia in neuronal mitochondria, Glutamate excitotoxicity, reduced availability of acetylcholine in the synaptic cleft, lymphoid tissue formation and antigenic variability in neuronal subpopulations. (Magliozzi, Howell et al. 2007, Geurts and Barkhof 2008, Trapp and Stys 2009, Klaver, De Vries et al. 2013, Stassart, Möbius et al. 2018) In addition, failure to control latent EBV infection in an immune privileged site, like the subarachnoid space could possibly result in recurrent intrathecal EBV reactivation and tissue damage in the surrounding GM regions. (Angelini, Serafini et al. 2013, Lossius, Johansen et al. 2014) Nevertheless, numerous other studies have been unsuccessful in detecting EBV in the MS brain lesions. (Willis, Stadelmann et al. 2009, Sargsyan, Shearer et al. 2010) and this remains to be an extremely debated controversy(Lassmann, Niedobitek et al. 2011).

Alternatively, there could be an involvement of an infectious agent in cortical demyelination and neuronal loss with primary tropism for oligodendrocytes and/or cortical neurons(Borkosky, Whitley et al. 2012), although no evidence exists yet for this likelihood in the disease. Further mechanisms targeting adaptive and/or innate immune responses to the GM include alterations, that are specifically linked to the function and/or degeneration of neurons, oligodendrocytes or astrocytes, such as metabolic changes(Zeis, Graumann et al. 2008), expression of post-translationally modified proteins and/or peptides(Mastronardi and Moscarello 2005), excitatory neurotransmitter release (Baranzini, Srinivasan et al. 2010), changes in electrical activity and/or ion currents and cytokine and/or cytokine receptor expression(Neumann, Cavalié et al. 1995).

### Regional GM atrophy in MS

The thalamus has cortical, subcortical and cerebellar connections. Hence, it is known to be a critical node in networks that support cognitive functions, memory processes, executive functions including information processing and attention. (Fama and Sullivan 2015)

Recent imaging as well as pathology studies have demonstrated the involvement of thalamus even in early MS. Atrophy of the thalamus has been observed in early RRMS(Trapp, Peterson et al. 1998), CIS at presentation(Štecková, Hluštík et al. 2014) and pediatric MS(Haider, Simeonidou et al. 2014). The volume loss in thalamus is possibly due in part to disconnection created by WM lesions(Azevedo, Overton et al. 2015). There is also evidence of a relationship between thalamic volume and WM lesion volume within thalmocortical projections, utilising diffusion tensor imaging tractography. (Lassmann 2014)

Cifelli and colleagues(Cifelli, Arridge et al. 2002) conducted a study of normalised thalamic volume (NTV) measurements in SPMS patients. Volumes of manually outlined thalami were normalised by intracranial volumes and showed a mean decrease of 17%. A moderate correlation was also observed third ventricular enlargement. According to the study conducted by Houtchens and colleagues(Houtchens, Benedict et al. 2007), the enlargement of third ventricle indicates a strong correlation with cognitive impairment in MS, indicating the clinical relevance of damage caused to the surrounding structures, such as the thalamus. Furthermore, the normalised thalamic volume was found to be 16.8% lower in the MS group as compared to healthy controls.

ClusterZ found a cluster in the left putamen with significant mean effect size of −0.75. The putamen is a part of the dorsal striatum and the basal ganglia and, plays a role in the regulation of movement, coordination, motor function and cognition. (Henry, Shieh et al. 2008, Batista, Zivadinov et al. 2012, Modica, Zivadinov et al. 2015) It is also involved in modulation of sensory and motor aspects of pain. (Munakomi.) Thus, a pathology like, neurodegeneration, might be expected to cause a broad spectrum of clinical manifestation from motor dysfunction to variable psychiatric disorder. (Koikkalainen, Hirvonen et al. 2007, Uono, Sato et al. 2017)

There is involvement of the left putamen in language functions such as bilingual language processing (Abutalebi, Della Rosa et al. 2012) and production, with some authors debating about the functional segregation of anterior and posterior putamen(Oberhuber, Parker Jones et al. 2013).

Employing FMRIB’s Integrated Registration & Segmentation Tool (FIRST)(Patenaude, Smith et al. 2011) as a tool for segmentation, previous researches have demonstrated progressive atrophy of the putamen in RRMS and SPMS(Bergsland, Horakova et al. 2012). Kramer and colleagues(Kramer, Meuth et al. 2015) recently showed putamen atrophy and its longitudinal progress during the disease course in MS. The quantitative analysis of putamen volume relative to disease duration showed a significant reduction in MS patients as compared to healthy controls. However, our study did not find a correlation between putamen atrophy and MS disease duration.

ClusterZ identified clusters in the right and left postcentral gyri with significant mean effect sizes of −0.74 and −0.66 respectively for GM atrophy in MS or CIS. BA3 is an eminent gyrus in the primary somatosensory cortex that is the sensory receptive area for the sense of touch. Li et al(Li, Jewells et al. 2013) used diffusion tensor imaging (DTI) and demonstrated neuroconnectivity changes in the left postcentral gyrus and the putamen, along with other regions. The left postcentral gyrus showed reduced communicability correlating with the 25-foot walk test results. Han and colleagues(Han, Tian et al. 2017) performed VBM analysis that revealed significant difference in GM between MS patients and controls. This decreased GM was observed in the right temporal lobe (caudate nucleus), right parietal lobe (postcentral gyrus) and right insula, along with other gyri, of RRMS patients which is partially consistent with our results.

ClusterZ revealed clusters in the left superior temporal gyrus with significant mean effect size of - 0.68. The superior temporal gyrus is associated with auditory and speech comprehension (Leff, Schofield et al. 2009, Friederici 2012) and perception of emotions in facial stimuli(Bigler, Mortensen et al. 2007, Radua, Phillips et al. 2010). In addition, it is an essential structure in the pathway containing prefrontal cortex and amygdala that are responsible for social cognition processes. (Bigler, Mortensen et al. 2007, Michl, Meindl et al. 2014) The study conducted by Achiron and colleagues (Achiron, Chapman et al. 2013), which compared RRMS patients and healthy controls, suggested correlation between reduced cortical thickness in superior temporal gyrus and global cognitive score, attention, information processing speed and motor skills.

ClusterZ revealed a cluster in the right insula with a significant effect size of −0.58. Following the removal of the temporal lobe for treating drug-refractory seizures in epileptic patients, stimulation of the exposed inferior part of the insular cortex provoked a variety of visceral sensory and motor responses, as well as somatic sensory responses, especially in the face, tongue, and upper limbs. This underwrote the conception of the insula as a primarily visceral-somatic region. (Uddin, Nomi et al. 2017) Studies have shown relation between functional connections of the basal ganglia and insula and fatigue severity in case of MS patients. (Finke, Schlichting et al. 2014, Jaeger, Paul et al. 2018, Lin, Zivadinov et al. 2018)

ClusterZ revealed a cluster in the right caudate body with significant mean effect size of −0.57. The mammalian basal ganglia structures, specifically the dorsal striatum, have been considered to have a role in memory and learning. (Packard and Knowlton 2002)

### Correlations of GM atrophy with clinical variables

The occurrence of brain tissue loss is not quite uniform, being more evident in brain GM in case of progressive MS, affecting some cortical and DGM regions more than others (Bendfeldt, Hofstetter et al. 2012). In vivo MRI-clinical correlation studies have identified significant associations of GM atrophy with cognitive impairment, physical disability and progressive multiple sclerosis that are not dependent on other imaging abnormalities, such as white matter lesion load. All-in-all, there are compelling reasons to try to better understand the mechanisms of grey matter atrophy and the subsequent neurodegeneration that is reflected. (Chard and Miller 2016)

Thalamic volume is a candidate MRI-based marker associated with neurodegeneration that can be utilized to hasten development of neuroprotective treatments. According to the research conducted by Azevedo and coworkers utilizing FreeSurfer longitudinal pipeline, decline in thalamic volume was observed to be significantly quicker in MS subjects compared to healthy controls(HC), with an estimated decline of −0.71% per year (95% confidence interval [CI] = −0.77% to −0.64%) in MS subjects and −0.28% per year (95% CI = −0.58% to 0.02%) in HC (p for difference = 0.007). The rate of decline was consistent throughout the MS disease duration and across MS clinical subtypes. Hence, thalamic atrophy was observed in early and consistently throughout the MS disease course, which is in agreement with the results of this study demonstrating correlation of disease duration with GM atrophy. (Azevedo, Cen et al. 2018) The authors pointed towards the deceleration of MS-specific brain volume change and the acceleration of age-related brain volume change during aging being more pronounced in the thalamus as compared to the whole brain, however it was not detectable in the putamen and caudate nuclei.

Datta and colleagues(Datta, Staewen et al. 2015) employed an automated pipeline based on tensor-based morphometry (TBM) to analyse GM atrophy in 1008 RRMS patients. Analysis of covariance was used to analyse atrophy differences between MS patients and HC on a voxel-by-voxel basis. Regional GM atrophy was observed in caudate, thalamus, putamen and cortical GM. Pearson’s correlation coefficient was used to analyse correlations between regional GM volumes and EDSS scores, disease duration and other variables. Weak correlations between thalamic volume and EDSS (r=-0.133; p<0.001) and DD (r=-0.098; p=0.003) were observed.

Significantly lower cortical and subcortical volumes have been reported in the insula, caudate, thalamus, putamen, amygdala and hippocampus by a cross-sectional VBM analysis of 3T T1-weighted MRI conducted by Palotai et al. (Palotai, Nazeri et al. 2019), adjusted for age, sex, disease duration and EDSS.

The present CBMA study results support these previous observations, taking into consideration the significant correlation between GM atrophy in the bilateral thalamus and MS disease duration.

### Previous meta-analyses conducted

A similar coordinate based meta-analysis in MS was conducted at the same time as the present study (Chiang, Wang et al. 2019). That study used the popular activation likelihood estimate (ALE) (Turkeltaub, Eden et al. 2002, Laird, Fox et al. 2005, Eickhoff, Laird et al. 2009, Turkeltaub, Eickhoff et al. 2012) algorithm. There are some limitations to the ALE algorithm that warrants consideration of the presented analysis using ClusterZ. Firstly, the ALE algorithm cannot account for studies reporting no coordinates, while ClusterZ considers these through censoring; ignoring such studies biases the analysis to the most significant results. Furthermore, the ALE algorithm works voxel-wise rather than cluster-wise, making it difficult to assess significance in merged clusters such as the left and right thalamic cluster, which is separated by the ClusterZ algorithm. Finally, excluding the reported Z scores puts meta-regression beyond the ALE algorithm. Nevertheless, the two analyses have produced remarkably similar results.

### Limitations

Even a well-performed meta-analysis of badly designed studies is bound to yield results that may contain inaccuracies. Bias and methodological limitations in the primary studies might be reflected in the meta-analysis. In the case of imaging studies methodological problems such as the use of uncorrected p-values are common. Coordinate based meta-analysis selects those results from the primary studies that are most concordant across the studies, and therefore goes some way to mitigate these problems. Nevertheless, CBMA results should be considered hypothesis generating in general and used to inform design of robust prospective studies. To this end the presented ClusterZ results provide a-priori regions of interest for testing as well as statistical effect size estimation for sample size calculations.

### Summary

GM atrophy regions involving the dorsal striatum, primary somatosensory cortex, insular cortex, auditory cortex and the relay station of the brain were identified cortically and subcortically in this meta-analysis of VBM studies of multiple sclerosis. The regions demonstrating significant effect sizes involved cortical and DGM regions, namely-bilateral Thalamus, bilateral BA3, Putamen, Caudate, BA13 and Superior temporal gyrus. Furthermore, disease duration was found to be a significant correlate of the standardized reported effect sizes in the thalamic clusters. These regions and the statistical effect size distributions can be used to design robust prospective studies of GM atrophy in MS.

## Supporting information

Primary analysis

Subanalysis (BMS)

Subanalysis (CIS)

Subanalysis (PPMS)

Subanalysis (RRMS)

Subanalysis (SPMS)

Covariate (age)

Covariate (disease duration)

Covariate (EDSS)

Forest plot for each significant cluster

Clinical characteristics of participants

DMTs for each included study

