## Supplementary figures and images for "Gray Matter alterations in MS and CIS: a Coordinate based Meta-analysis and regression"

### Forest plot for each significant cluster

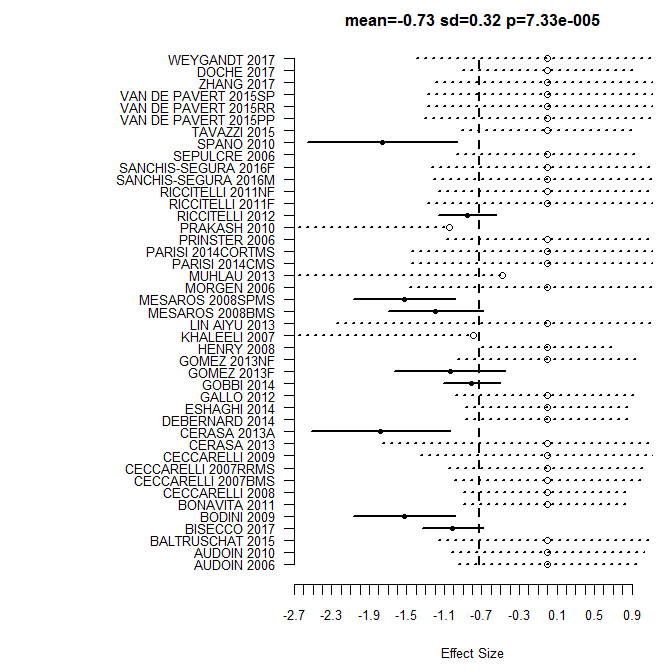

**4**

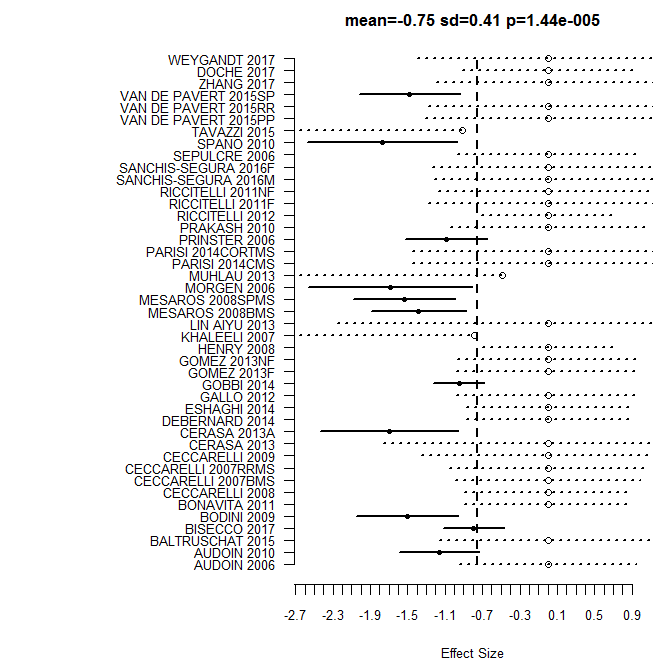

**3**

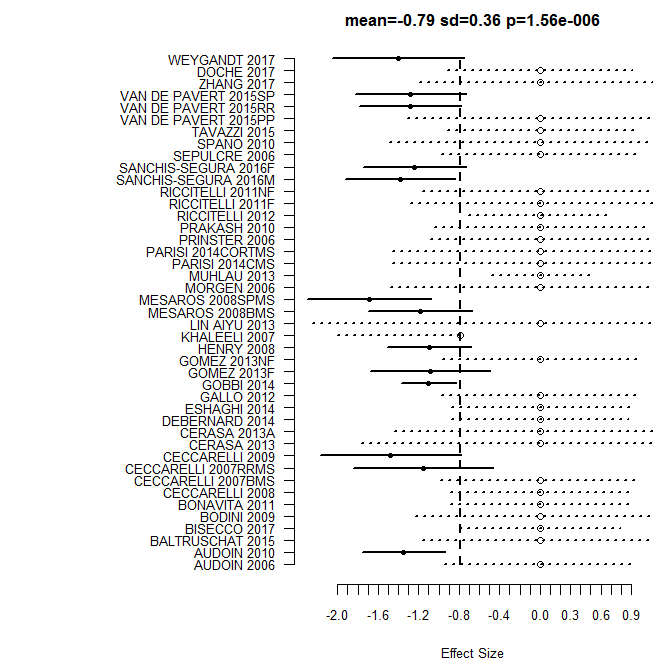

**6**

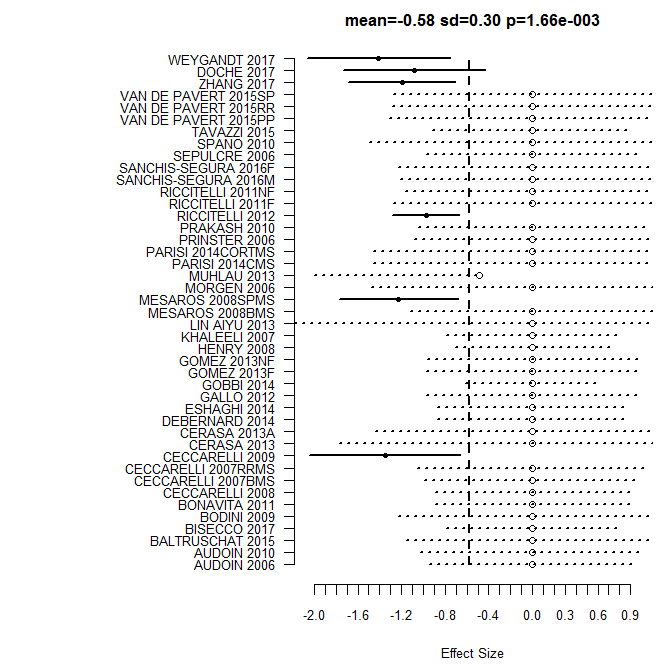

**5**

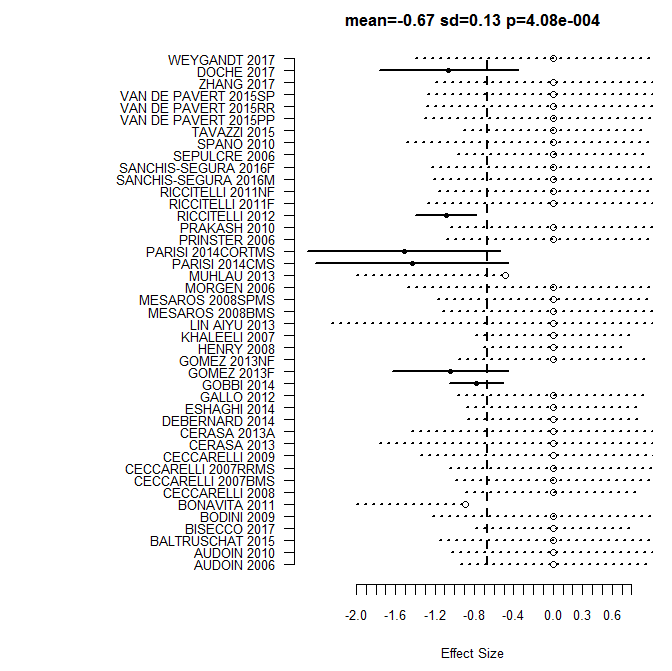

**8**

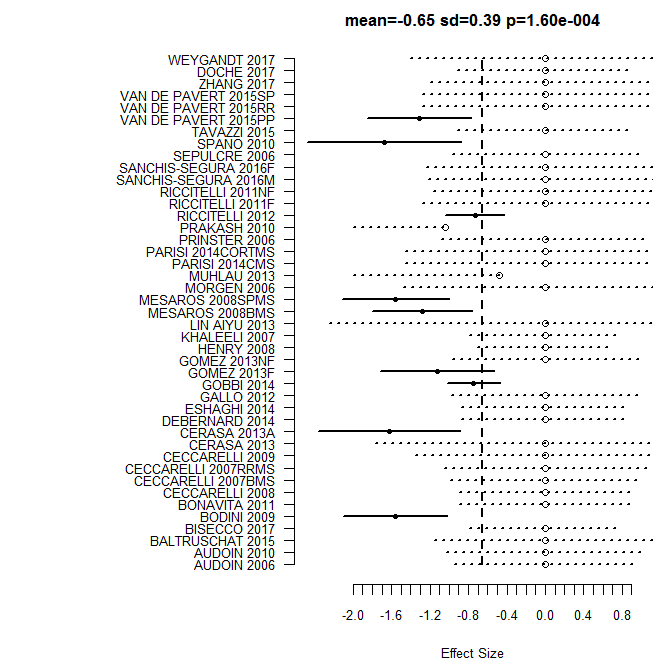

**7**
